# Population receptive field properties of the human visual claustrum zone

**DOI:** 10.64898/2026.02.02.703259

**Authors:** Carmen Anna Pizka, David Linhardt, Adam Coates, Dominik Zuschlag, Christian Windischberger, Natalia Zaretskaya

## Abstract

The claustrum is a thin, bilateral sheet of grey matter between the insula and putamen that stands out by its high interconnectivity with almost the entire cortex. Despite continuing research in humans and animals, its functional role remains largely unknown. In the present study, we explored the topographic organization of the recently described human visual claustrum zone. We performed a population receptive field (pRF) analysis on the 7T retinotopy dataset of the Young Adult Human Connectome Project (N = 181, 109 female) comparing the visual claustrum with established visual field properties of the lateral geniculate nucleus, the primary visual cortex, and higher-level topographic maps of the dorsal and the ventral stream. Our results demonstrate for the first time that the human visual claustrum showed several topographic properties typical for visual areas, including a representational bias towards the contralateral visual field, and a pRF size increase with increasing eccentricity. At the same time, the claustrum also exhibited a positive eccentricity gradient along the posterior-anterior axis, an extended representation of the visual periphery compared to other areas and a lack of horizontal meridian bias. These latter two properties highlight the claustrum’s role as a higher-level nucleus which is less dependent on sensory input. This study is the first to characterize the topographic organization of the visual claustrum zone in humans, highlighting its uniqueness among the known visually responsive regions.

**Significance Statement:** The claustrum is a thin subcortical brain region whose function is still largely unknown. Previous animal studies showing unimodal sensory zones within the nucleus suggest an involvement in sensory processing. The present study focuses on the visual claustrum zone and is the first to characterize its topographic organization in humans. We used population receptive field mapping for functional Magnetic Resonance Imaging on the 7T retinotopy dataset of the human connectome project. We could demonstrate similarities to other visually responsive areas and characteristics that distinguish the visual claustrum. Our study yields important information on the claustrum’s visuotopic organization that can guide further investigation of its role in visual processing and cognition.

## Introduction

Multiple subcortical nuclei in the brain are known to contribute to the processing of visual information. The claustrum, a bilateral sheet of grey matter concealed laterally by the insula and medially by the putamen, is only rarely acknowledged in this regard. Despite its fine anatomy with down to less than 1 mm lateral extent, it stands out by its highly networked nature as the most densely connected structure per regional volume with reciprocal connections to virtually all cortical and multiple subcortical regions (Kapakin, 2011; Milardi et al., 2015; Torgerson et al., 2015). As this profound connectivity suggests an involvement in higher-level cognition (Jackson et al., 2020), various hypotheses regarding the claustrum’s role have been put forward, linking it most prominently to the formation of conscious experience (Crick & Koch, 2005; Liaw & Augustine, 2023) as well as to attentional processing, pain perception, sleep control and hallucinations (Nichols, 2016; Atilgan et al., 2022). However, since its enclosed location and structural properties pose challenges for detailed investigation, the function of the claustrum remains largely unknown (Goll et al., 2015; Smith et al., 2020).

A well-established aspect of the claustrum’s physiology speaks for its involvement in perceptual processing. Several electrophysiological studies in animal models demonstrated unimodal visual, auditory, and somatosensory zones within the nucleus (e.g., Olson & Graybiel, 1980; Remedios et al., 2010). With respect to the visual domain, previous research in felines suggests a discrete visual zone in dorsocaudal portions of the claustrum, which exhibits orderly retinotopic connections to the striate and extrastriate cortices (Olson & Graybiel, 1980; LeVay & Sherk, 1981a,b). In contrast, studies in macaques indicate either a single or two separate visual zones in more ventrocaudal and central segments of the nucleus, reciprocally connected to visual cortices, but displaying merely a coarse topography (Remedios et al., 2010; Gattass et al., 2014). Diffusion-Weighted Imaging-based tractography studies imply that similar sensory zones may exist in humans (Milardi et al. 2015; Torgerson et al., 2015). Topographically organized connections were demonstrated with the superior frontal, precentral, postcentral, and posterior parietal cortex (Fernández-Miranda et al., 2008). Further, a recent functional Magnetic Resonance Imaging (fMRI) study using naturalistic video stimuli found visually evoked responses situated in a distinct ventral part of the nucleus, which is thought to be the human homologue of the visual claustrum zone (Coates et al., 2024; Figure 1A). Revealing the precise functional properties of this specialized sensory zone in humans and comparing it to animal findings will advance our understanding of the claustrum’s contribution to sensory processing.

**Figure 1.**
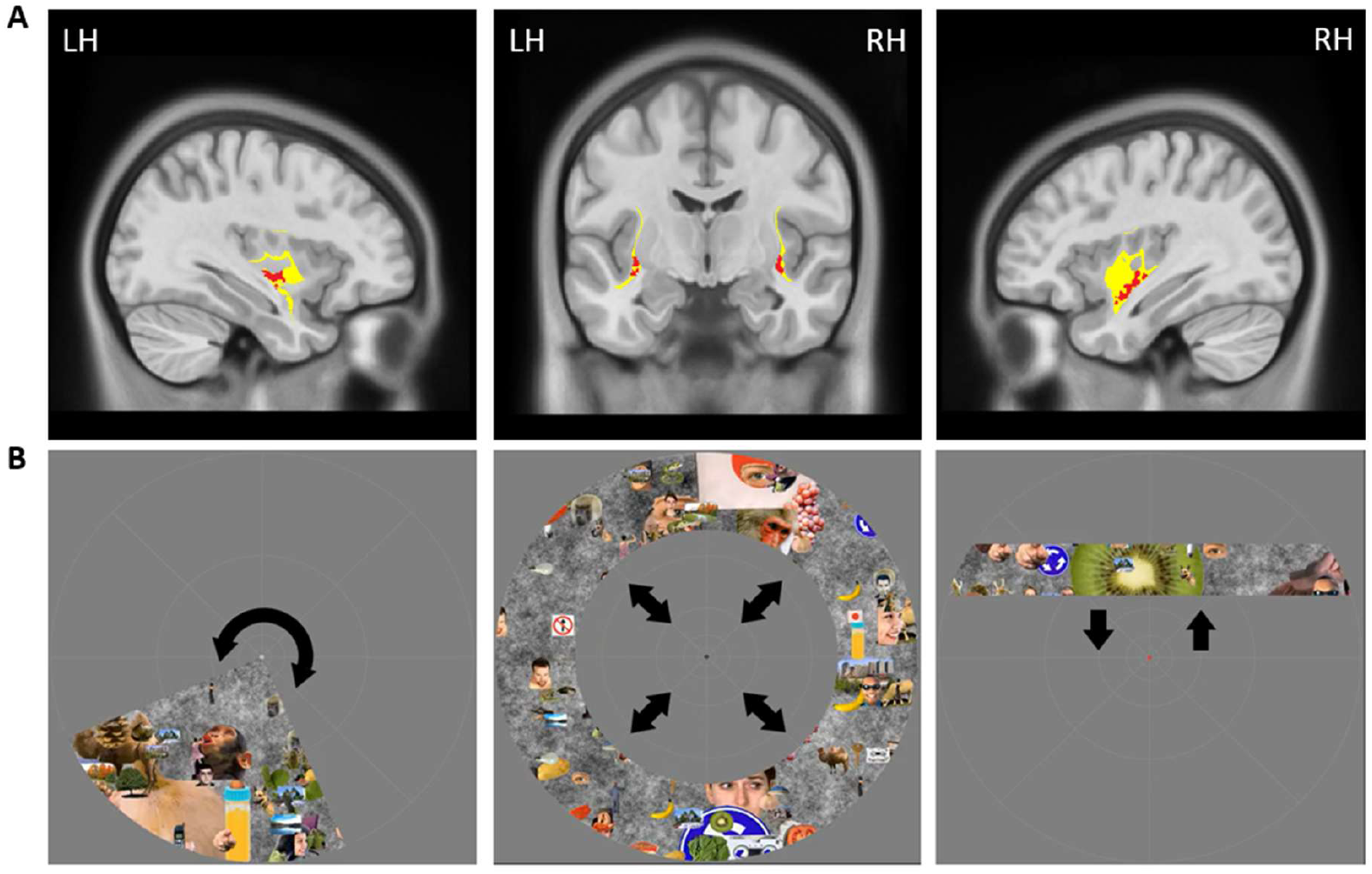
Illustration of the visual claustrum zone and the pRF mapping stimuli. A) Location of the visual claustrum zone (red) within the whole claustrum (yellow) depicted on the MNI152NLin2009cAsym-space template (mni_icbm152_t2_tal_nlin_asym_09c), with the left claustrum in sagittal view (left), the left and the right claustrum in coronal view (middle) and the right claustrum in sagittal view (right; RH: right hemisphere; LH: left hemisphere). B) The pRF mapping stimuli used by Benson et al. (2018) with the wedge (left), ring (middle), and bar (right) aperture uncovering the colorful object textures, and the color changing dot for the central attention task. Black arrows indicate the movement directions of the apertures. Horizontally and diagonally moving bar stimuli are not depicted.

A method that allows to characterize the topographic organization of cortical and subcortical visual areas using fMRI is population receptive field (pRF) mapping (Dumoulin & Wandell, 2008). The pRF method estimates the aggregated receptive field of the population of neurons contained within an individual voxel using a model-driven forward-encoding approach. Each voxel’s pRF is modelled as a two-dimensional Gaussian function with parameters for the center-position (x, y) and pRF size (σ), which are optimized in a voxel-wise fitting process by approximating the time course predictions to the obtained fMRI signal changes. Allowing for a more accurate (Dumoulin & Wandell, 2008) and comprehensive understanding of the topographic organization than conventional phase-encoded retinotopic mapping (Engel, 1997), the visual field maps obtained with the pRF approach indicate the location of each voxel’s pRF, including standard polar angle and eccentricity measures, as well as its spatial extent.

In the present study we explored the pRF characteristics of the human visual claustrum zone (VisCl) using the 7T retinotopy dataset of the Young Adult Human Connectome Project (HCP; Benson et al., 2018). We provide detailed comparisons with established pRF properties of the lateral geniculate nucleus (LGN), the primary visual cortex (V1), as well as two sets of higher-level topographic maps along the dorsal (intraparietal sulcus, IPS) and the ventral (ventral occipital, VO) streams, and find both similarities with other visual areas as well as unique claustrum characteristics.

## Materials and Methods

### Subjects

The HCP 7T retinotopy dataset of the 1200 release from 2017 includes the complete retinotopy data for a total of 181 subjects (age range 22–35, 109 female) that were used for statistical analysis in the present study. The dataset additionally includes the average time series across subjects (Sub-ID: 999999) which was used for a single visualization in the present study (Figure 6A). Participation criteria included eligibility for MRI examination and normal or corrected-to-normal visual acuity. For a detailed description of the sample, please see Benson et al. (2018).

### Procedure and stimuli

An essential feature of the HCP retinotopy study (Benson et al., 2018) is the use of colorful, dynamically changing real-life objects as part of the stimulus design. This design effectively drives neural responses in higher- and lower-level visual areas (Benson et al., 2018). For visual stimulation the standard apertures for pRF mapping were employed, including clockwise and counter-clockwise rotating wedges, expanding and contracting rings, and bars that moved vertically, horizontally, and diagonally across the screen. These apertures uncovered colorful changing object textures. A total of 100 distinct textures were generated by overlaying 92 different sized chromatic images of various real objects, such as faces, animals, and buildings, on an achromatic pink-noise background using random placement. During presentation, the dynamic apertures were updated at a rate of 15 Hz with one of the texture images being randomly selected on each aperture update without being shown consecutively. Apertures and textures covered a circular area of the visual field of approximately 16° of visual angle in diameter, with a uniform grey display beyond that region. Benson et al. (2018) suggest that the vivid object textures induce higher blood oxygenation level-dependent (BOLD) responses and enhance test-retest reliability of retinotopic estimates compared to conventional black-and-white reversing checkerboard patterns. As the visual claustrum zone is known to respond well to naturalistic stimuli in primates (Remedios et al., 2010) and humans (Coates et al., 2024), this special stimulus design makes the dataset ideally suited for investigating visual responses of the claustrum.

To support fixation and allocation of attention to the center of the display, a central attention task was presented. Subjects had to respond to a dot (0.38 × 0.38°), changing color between black, white, and red every 1-5 s, via button press (Current Designs, Philadelphia, PA, USA). Additionally, a semi-transparent fixation grid was superimposed on the display to further encourage a stable gaze focus (Benson et al., 2018). A NEC NP4000 projector with a resolution of 1024 × 768 at 60 Hz was used to present the stimuli on a back-projection screen, which the subjects could view from a distance of 101.5 cm through a mirror attached to the head coil. Examples for the presented stimuli are shown in Figure 1B.

### Imaging and preprocessing

Acquisition and preprocessing of structural and functional MRI data was performed by Benson et al. (2018) according to the HCP pipelines (Glasser et al., 2013; Vu et al., 2017). High resolution structural T1- and T2-weighted scans were obtained at 0.7 mm isotropic resolution using the HCP’s customized 3T system (Connectome Skyra; Siemens Healthineers, Erlangen, Bayern, Germany) with a 32-channel head coil (Van Essen et al., 2013). Preprocessing of structural data included cortical surface reconstruction (Glasser et al., 2013), a two-step surface-based registration to the HCP 32k fsLR standard space using cortical-folding-based (MSMSulc) and areal-feature-based multimodal surface matching (MSMAll) driven partly by resting-state visuotopic maps to improve alignment of pRF solutions (Robinson et al., 2014, 2018; Glasser et al., 2016a), and nonlinear volume-based registration to the volumetric standard template MNI152NLin6Asym using FNIRT (Glasser et al., 2013).

Functional data were obtained with 1.6mm isotropic resolution at 7T (MAGNETOM actively shielded scanner; Siemens Healthineers, Erlangen, Bayern, Germany) using a 32-channel receiver coil array with a single channel transmit coil (Nova Medical, Wilmington, MA, USA). Preprocessing of functional data included head motion correction, EPI spatial distortion correction, registration of cortical data to the HCP 32k fsLR standard surface space using MSMAll (Robinson et al., 2014, 2018; Glasser et al., 2016a), denoising for spatially specific structured noise using multirun spatially independent component analysis plus FMRIB’s ICA-based X-noiseifier (sICA+FIX; Griffanti et al., 2014; Salimi-Khorshidi et al., 2014; Glasser et al., 2018) and nonlinear registration of volumetric data to MNI152NLin6Asym using FNIRT with spline interpolation to minimize blurring effects (Glasser et al., 2016b; Coalson et al., 2018). The data produced by the HCP preprocessing pipelines come in varying spatial resolution. For the present study, only the 1.6 mm functional data in MNI152NLin6Asym space were used.

The preprocessed data of the 1200 release from 2017, which were used for this study, were formerly available from ConnectomeDB (https://www.humanconnectome.org/study/hcp-young-adult). An updated form of the dataset that underwent different preprocessing is currently available with registration from ConnectomeDB powered by BALSA (https://balsa.wustl.edu/).

### Definition of regions of interest

#### Visual claustrum zone

Voxels corresponding to the visual claustrum zone were derived from a high-resolution 7T fMRI study in 15 healthy subjects, which identified the region of the visual claustrum in humans (Coates et al, 2024). The mask was defined as voxels that fall within the whole anatomical claustrum mask (Coates & Zaretskaya, 2024) and exhibit visual activity at a cluster forming threshold of p <.01 (uncorrected) and a cluster significance level of p <.05 (FWE-corrected) at the group level.

Since the functional HCP data were coregistered to the volumetric MNI152NLin6Asym, the visual claustrum mask was mapped from its original MNI152NLin2009cAsym space to MNI152NLin6Asym to ensure compatibility. We used ANTS symmetric normalization algorithm (antsRegistrationSyN.sh; Avants et al., 2008; Klein et al., 2009) with rigid, affine and nonlinear, diffeomorphic registration on the standard templates of the respective MNI spaces. The visual claustrum mask was down-sampled from 0.75 mm to 1 mm to match the resolution of the templates. Then the affine transformation matrix and the nonlinear deformation field generated during template registration was applied to the visual claustrum mask using antsApplyTransforms with linear interpolation. Afterwards, the visual claustrum mask was down-sampled to 1.6 mm, cropped by one side slice, to match the resolution and dimension of the functional HCP data, and binarized.

#### Comparison regions

Several visual regions with well-characterized pRF properties were defined for comparison. The LGN was defined using the probabilistic MNI152NLin6Asym-space atlas based on microstructure-sensitive high-resolution 7T in-vivo MRI assessments of 27 healthy subjects and post-mortem analysis of a human LGN specimen in combination with histological validation methods (Müller-Axt et al., 2021). To match the resolution and dimension of the functional HCP data, the LGN mask was resampled from its original 1 mm to 1.6 mm isotropic voxel size and binarized. For the V1, IPS, and VO corresponding maximum probability maps (V1v-V1d, IPS0-IPS4, VO1-VO2) were taken from the topographic MNI152NLin6Asym-space atlas by Wang et al. (2015) and are based on 3T fMRI task data of 31 to 50 subjects.

### GEM-population receptive field mapping

For pRF mapping we followed the fast GPU-empowered (GEM) approach by Mittal et al. (2026), which is based on the standard method by Dumoulin and Wandell (2008). We estimated pRF’s using a 2-D Gaussian model defined by visual-field position (x, y) and pRF size (σ). For the HCP dataset (six runs), we generated two independent parameter sets: one from the concatenated bar-aperture runs and one from the concatenated wedge and ring runs. The binarized aperture sequence was convolved with a double-gamma HRF, identical across subjects and regions. Model time courses were generated by multiplying the convolved stimulus with the 2-D Gaussian pRF and integrating over the visual field. pRF estimation proceeded in two stages. First, coarse parameters were obtained using a full grid search within the predefined sampling space. Second, these parameters were refined by assuming that the local error landscape around the coarse-fit point is smooth and can be approximated by a quadratic function, enabling a non-iterative update. Final pRF centers (x, y) were obtained in Cartesian coordinates and converted to polar coordinates to compute standard polar angle and eccentricity metrics. Since the goodness-of-fit for the pRF models is expected to decrease in higher-level visual areas (Szinte & Knapen, 2020) and in subcortical structures due to partial volume effects and lower SNR (DeSimone et al., 2015), the reported results are based on a variance explained threshold of 1.5% for all ROIs.

Noteworthy, the type of employed aperture can bias pRF mapping results, including the spatial distribution of pRF centers, the goodness-of-fit, and the estimated pRF size and eccentricity values (Alvarez et al., 2015; Linhardt et al., 2021; Pawloff et al., 2023). For central visual field regions, wedge/ring apertures contribute to a higher pRF center density due to higher variance explained values compared to bar apertures. In more peripheral field regions, stimulation with bar apertures produces a more homogenous coverage of pRF centers. Further, pRF size and eccentricity estimates are generally reduced by wedge/ring apertures compared to bar stimuli. To address these aperture-dependent biases, we combined the retinotopy data of the concatenated wedge and ring aperture runs with the data of the concatenated bar aperture runs. The combination was based on the variance explained metric of each aperture type model, adopting a Best-of-Two strategy (Linhardt et al., 2021). For each voxel, the pRF parameters were chosen from the aperture type whose prediction yielded a higher variance explained. As a result, every voxel is represented once in the data and its corresponding pRF parameters are derived from only one of the two aperture type models.

### Experimental design and statistical analysis

For the characterization of pRF properties of the human visual claustrum, we quantified several major visual field representation properties described in detail below. All measures were calculated per subject, hemisphere and ROI, with subsequent parametric statistical inference where applicable. For all analyses, the pRF mapping parameters were filtered for physiologically implausible values. As Benson et al. (2018) note that their reported stimulus size of 16° in diameter should be considered approximate due to variations in the setup, pRF size was filtered for values > 18°, eccentricity for values > 9°. Furthermore, since the smallest pRF size values commonly reported for visual regions such as V1, V2 or V3 corresponds to around 0.5°-2° (Dumoulin & Wandell, 2008), we additionally filtered pRF size for values ≤ 0.3°. Statistical tests were always two-tailed. All analyses were carried out in a Jupyter Notebook environment with Python (Version 3.12.11) using the packages NumPy, SciPy, Pingouin, Pandas, Matplotlib and Seaborn.

#### Code Accessibility

The GEM-pRF software package used for the pRF analysis is publicly available via PyPI (https://pypi.org/project/gemprf/). The custom Python scripts used for analysis in the present study are publicly available on the Open Science Framework (the URL will be included here).

#### Lateralization bias

Lateralization of the visual field representation was quantified as the average voxel count across subjects for the pRF x-position per 1° bin and examined by plotting histograms for each hemisphere. For the purpose of statistical inference, we calculated the average x-position across voxels per hemisphere and tested the difference in mean horizontal field position between left and right hemisphere for each ROI using a paired t-test. Additionally, we also quantified the laterality index across hemispheres per ROI, by calculating the difference between the number of ipsi- and contralaterally responsive voxels relative to the total number of voxels. Left and right hemisphere ROI data were pooled together for this analysis. The resulting index ranges from −1 (100% contralateral) to +1 (100% ipsilateral), passing through 0 (50% contralateral and 50% ipsilateral). We tested the differences in average laterality index between the claustrum and each of the four other ROIs using paired t-tests with Bonferroni correction (α_corr_ = 0.0125) to account for multiple comparisons.

#### Meridian representation

Since many visual areas exhibit a bias towards representing the horizontal meridian (Schneider et al., 2004; DeSimone et al., 2015; Benson et al., 2021; Himmelberg et al., 2021), we tested for the presence of such bias in the visual claustrum. We examined the average fractional volume across subjects as a function of polar angle. We plotted circular histograms for each hemisphere, showing 20 radial segments, each spanning 18° for visualization. To further quantify the extent of horizontal and vertical meridian representation, we pooled the data over both hemispheres and counted the voxels that exhibited polar angle values within wedges with a width of ± 45° centered either on the left and right horizontal meridian or the upper and lower vertical meridian. We then tested for voxel count differences between the horizontal and the vertical field coverage using a paired t-test for each ROI. To compare the claustrum to the other areas, we calculated the meridian bias, defined as the difference in voxel count between the horizontal and the vertical field coverage, and performed pairwise comparisons between the claustrum and the other ROIs using paired t-tests with Bonferroni correction (α_corr_ = 0.0125).

#### pRF size increase with eccentricity

The systematic increase of pRF size with eccentricity, typical for visual areas (e.g., Dumoulin & Wandell, 2008; DeSimone et al., 2015), was analyzed by performing linear fits to the eccentricity versus pRF size data for each subject and then computing a one-sample t-test on the slopes against a population mean of zero. We compared the claustrum’s average slope parameter with the slope parameters of the other ROIs using paired t-tests with Bonferroni correction (α_corr_ = 0.0125).

#### Eccentricity bias

Given the preference for stimulation of the visual periphery reported for the feline visual claustrum zone (LeVay & Sherk, 1981b), we tested for the presence of this bias in our human data. To quantify the proportions of voxels with central and peripheral field representations, we examined the fractional volume as a function of eccentricity for each hemisphere by means of a histogram and calculated the eccentricity value at the 50% fractional volume threshold. Furthermore, we computed the average eccentricity value of both hemispheres per subject. We compared the average eccentricity values at 50% threshold between the claustrum and the other ROIs using paired t-tests with Bonferroni correction (α_corr_ = 0.0125). The same statistical analysis was performed for the average eccentricity value.

#### Relation between topography and anatomy

To determine if the claustrum’s topographic organization is ordered with respect to the brain’s gross anatomy, we pooled the data over both hemispheres and computed the linear fits for eccentricity as a function of the MNI space y- (posterior-anterior) and z- (inferior-superior) coordinate for each subject before testing for nonzero slopes using one-sample t-tests. For the MNI space x-coordinate (left-right) we analyzed the data for the right and the left hemisphere separately. Following the same approach, we also computed the linear fits for the polar angle as a function of the three MNI space coordinates for each subject. To be able to combine polar angle data of both hemispheres we transformed the polar angle values of each hemisphere to go from 0° (bottom) to 180° (top). We then again tested for nonzero slopes with one-sample t-tests.

## Results

Table 1 shows the total voxel count for each ROI, the average number of filtered voxels after thresholding by variance explained and combining the pRF data across bar and wedge/ring tasks, and the average number of analyzed voxels after excluding physiologically implausible values.

**Table 1.** Regions of Interest statistics. The total voxel count, the average fraction of filtered and combined voxels, and the average fraction of analyzed voxels across subjects for each ROI are shown per hemisphere. Note the hemispheric asymmetry in the total voxel count for the visual claustrum, with more voxels in the right than in the left hemisphere.

| Left hemisphere |  |  |  | Right hemisphere |  |  |
| --- | --- | --- | --- | --- | --- | --- |
|  | Total | R<sup>2</sup>-filtered combo | Analyzed | Total | R<sup>2</sup>-filtered combo | Analyzed |
| VCL | 61 | 36.28% | 12.72% | 118 | 37.81% | 12.83% |
| LGN | 121 | 57.01% | 31.16% | 111 | 58.43% | 31.62% |
| V1 | 1673 | 91.87% | 52.64% | 1467 | 91.07% | 52.24% |
| IPS | 897 | 83.44% | 56.73% | 973 | 81.84% | 56.07% |
| VO | 364 | 84.23% | 55.00% | 406 | 82.30% | 53.80% |

### Lateralization bias

The visual claustrum zone exhibited a representational bias towards the contralateral visual field (Figure 2A), as was the case in other visual areas examined (Figure 2B), and confirmed by statistical analysis comparing average x-position between hemispheres (Figure 2C; claustrum: t(179) = 4.67; p <.001; d = 0.35; all other areas: t(180) ≥ 37.05, p <.001, d ≥ 2.75). Correspondingly, the laterality indices for all inspected regions were negative, indicating a predominant representation of the contralateral hemifields (Figure 2D). However, the laterality index was significantly lower for the visual claustrum than for each of the other regions, indicating a weaker lateralization of visual responses. For all Bonferroni corrected paired t-tests comparing the claustrum with the other visual areas: t(180) ≥ 17.00, p_corr_ <.001, d ≥ 1.26.

**Figure 2.**
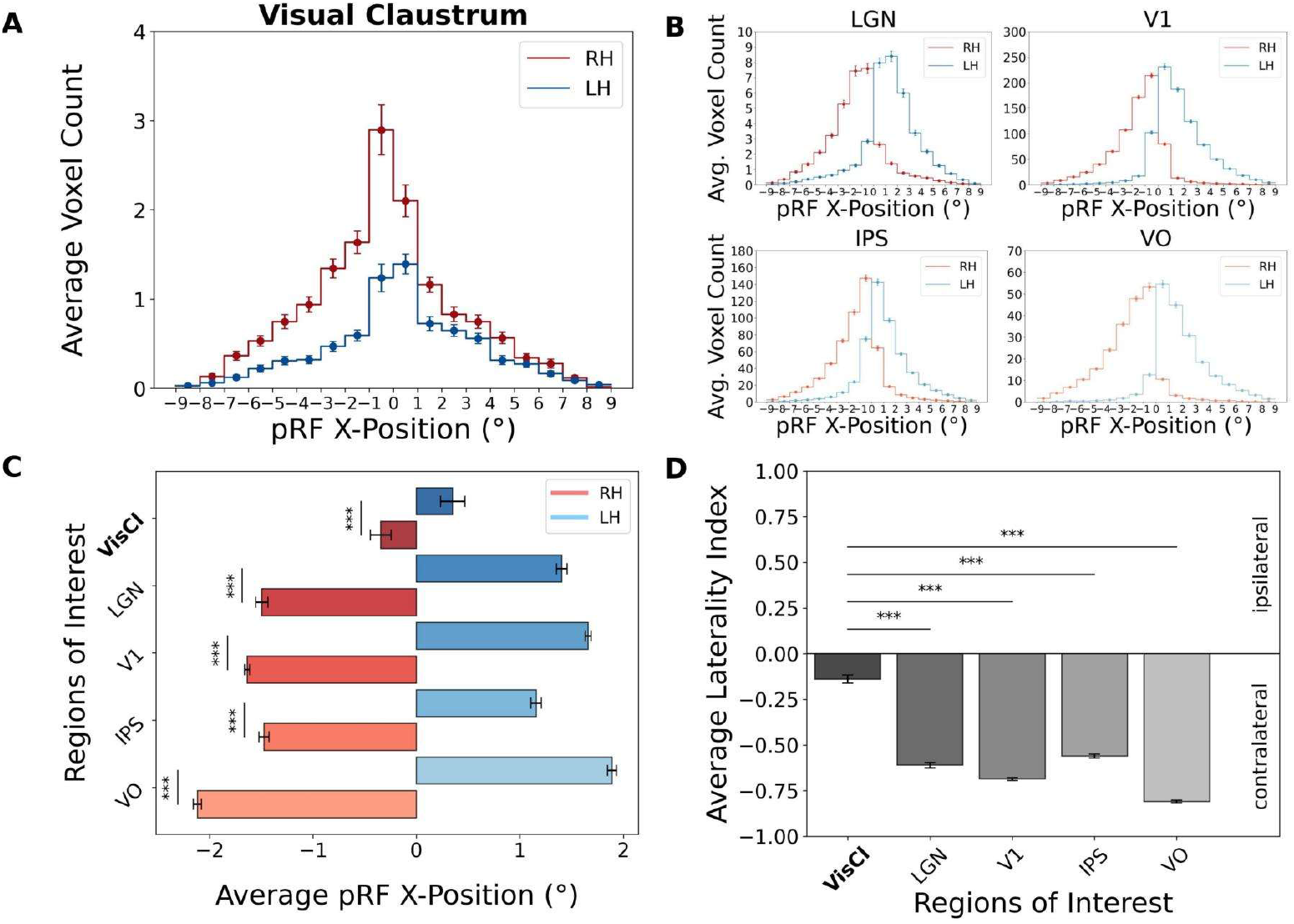
Contralateral visual field representation bias in the visual claustrum and other regions. Data for the right hemisphere (RH) is depicted in red and for the left hemisphere (LH) in blue. A-B) Histograms of the average voxel count across subjects for the pRF x-position per 1° bin for A) the visual claustrum zone and B) other examined areas. C) Average pRF x-position for each hemisphere in each ROI, demonstrating biases of each hemisphere towards the contralateral visual field. D) Laterality index for each ROI defined as the difference between the number of voxels with ipsilateral and contralateral responses, divided by the total number of voxels. Negative values correspond to the contralateral visual field bias. Error bars represent the SEM over subjects.

### Meridian representation

The claustrum showed a homogenous distribution of polar angles across visual space, with no significant difference in the extent of vertical and horizontal meridian representations (t(180) = 0.07, p =.948, d = 0.01), while the other regions exhibited an overrepresentation of the horizontal meridian and an underrepresentation of the vertical meridian (all other areas: t(180) ≥ 3.20, p ≤.002, d ≥ 0.24; Figure 3C). This effect is also reflected in the circular histogram plots for the visual claustrum (Figure 3A) and for the other areas (Figure 3B). The comparison of the meridian bias values between the claustrum and the other visual areas confirmed that the extent of the horizontal and vertical meridian representation in the claustrum differs significantly less than in the other ROIs (For all Bonferroni corrected paired t-tests: t(180) ≤ −3.20, p_corr_ ≤.007, d ≤ −0.24; Figure 3D).

**Figure 3.**
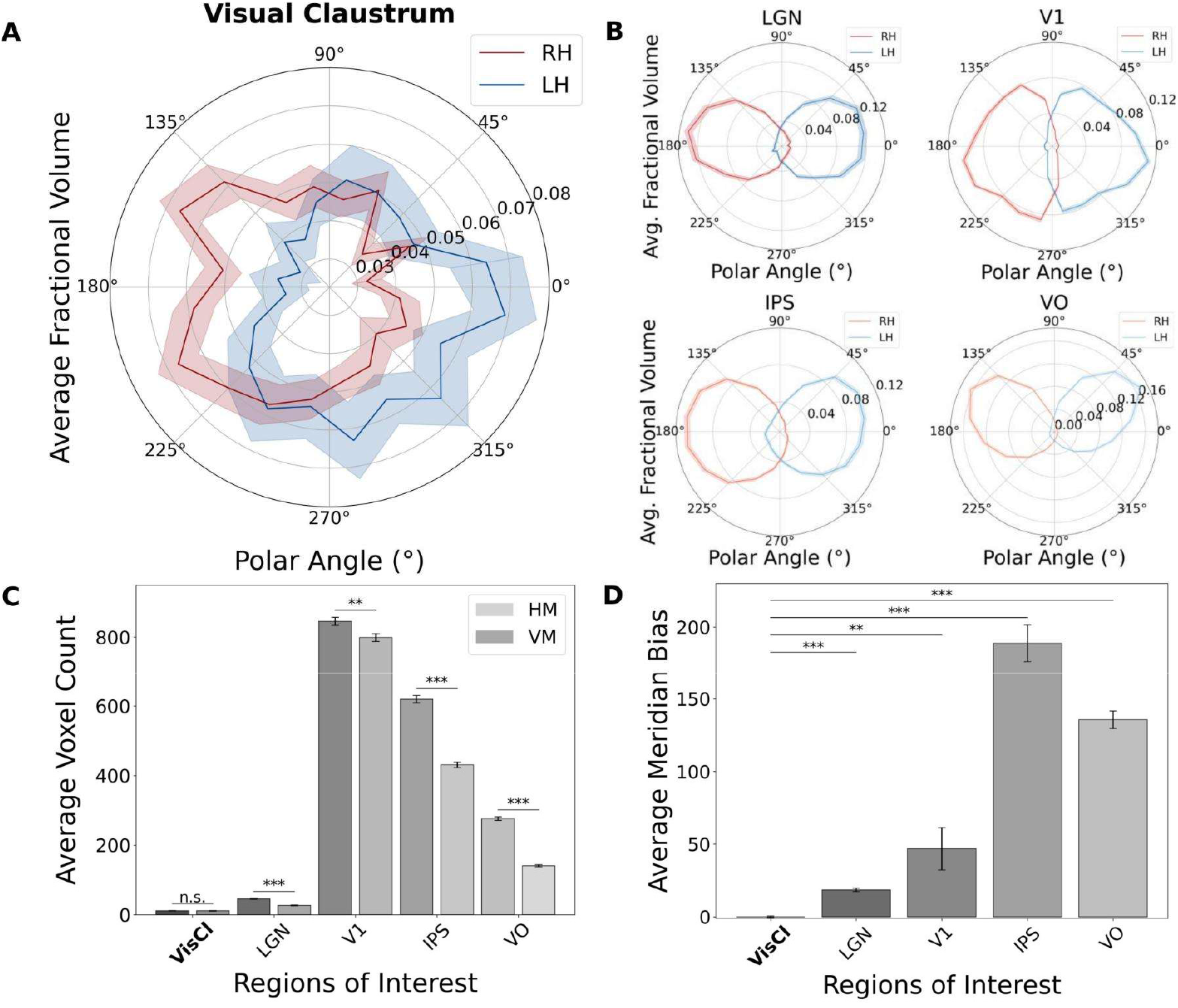
Distribution of polar angle values for the right (RH; red) and the left hemisphere (LH; blue) in A) the visual claustrum and B) other examined areas. Compared to other areas, which exhibit an overrepresentation of the horizontal meridian, the claustrum shows a homogenous distribution of polar angles values. C) Extent of horizontal (HM) and vertical meridian (VM) coverage for each ROI. D) Average meridian bias for each ROI. Shaded lines and error bars represent the SEM.

### pRF size increase with eccentricity

Similar to other examined visual regions, the visual claustrum showed a significant positive relationship between pRF size and eccentricity (t-test of the slope against zero for the claustrum: t(180) = 2.09; p =.038, d = 0.16; all other areas: t(180) ≥ 14.08, p <.001, d ≥ 1.05; Figure 4A shows the slope for the claustrum and 4B for the other ROIs in a single subject). This association between pRF size and eccentricity was significantly weaker in the claustrum compared to other ROIs (For all Bonferroni corrected paired t-tests: t(180) ≤ −5.19, p_corr_ <.001, d ≤ −0.39; Figure 4C).

**Figure 4.**
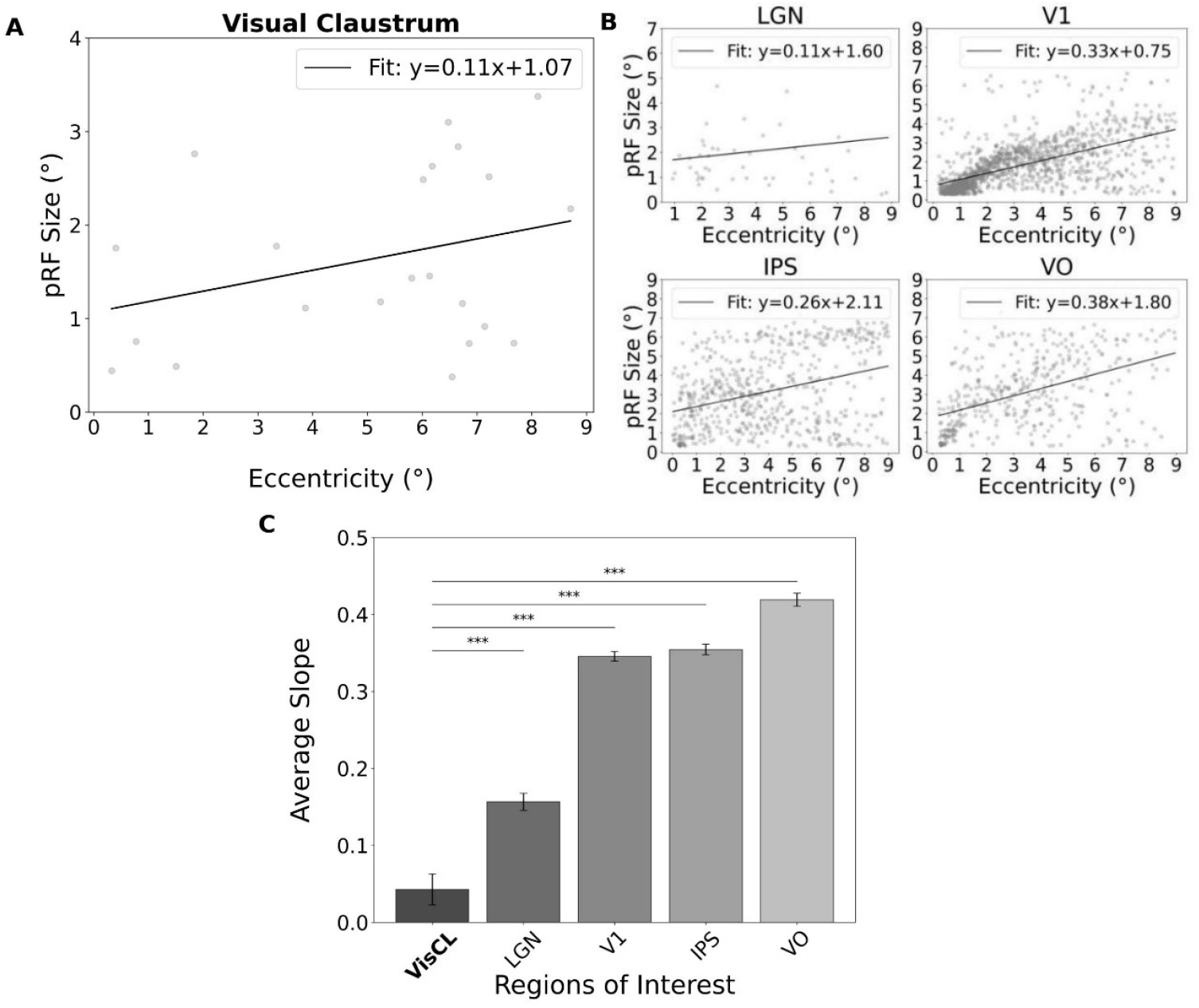
Relationship between eccentricity and pRF size for an example participant (Sub-ID: 159239) in A) the visual claustrum and B) other examined regions. C) Slope values of the regression line for each area, demonstrating a weaker but significant association between eccentricity and pRF size in the claustrum. Error bars represent the SEM.

### Eccentricity bias

Compared to other regions, the visual claustrum demonstrated a bias towards more peripheral visual field locations. Across subjects 50% of its voxels represented the central 3.67° of eccentricity in the visual field on average (Figure 5A). The other ROIs showed larger representations devoted to the central visual field, with 50% of their voxels representing the central 2.20°–2.81° (Figure 5B). Statistical comparison confirmed a significantly higher average eccentricity at 50% fractional volume threshold for the visual claustrum than for other areas (For all Bonferroni corrected paired t-tests: t(180) ≥ 9.95, p_corr_ <.001, d ≥ 0.74; Figure 5C). The tuning for more peripheral visual areas in the claustrum relative to the other regions was further confirmed by comparing the average eccentricity value in the visual claustrum with the values in the other areas (For all Bonferroni corrected paired t-tests: t(180) ≥ 10.83, p_corr_ <.001, d ≥ 0.81; Figure 5D).

**Figure 5.**
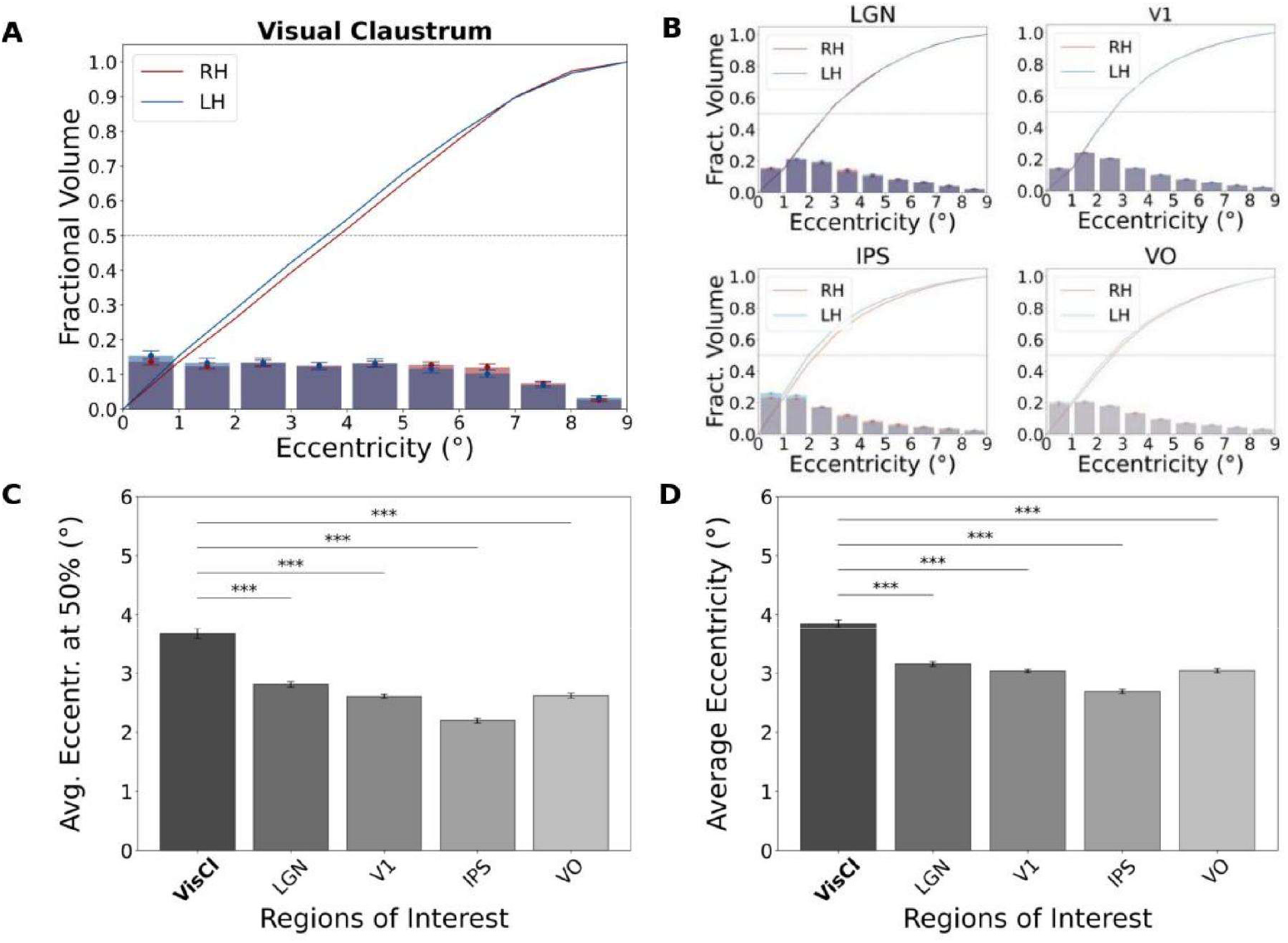
Eccentricity representation biases in the claustrum and other visual areas. A-B) The fractional volume as function of eccentricity for the right (RH; red) and the left hemisphere (LH; blue) of A) the visual claustrum zone and B) other examined areas. The bars represent the number of voxels within each 1° bin of eccentricity and the lines the cumulative sum across bins per hemisphere. C) Average eccentricity at 50% fractional volume for each ROI. D) Average eccentricity value for each ROI. Error bars represent the SEM.

### Relation between topography and anatomy

Finally, we checked if there is a systematic change of the eccentricity and polar angle values along the three brain axes, which would suggest the existence of an ordered retinotopic map. Eccentricity values increased significantly from posterior-to-anterior positions (Figure 6), with more anterior regions representing more peripheral visual field locations (*M* = 0.04, *SD* = 0.22, *t*(180) = 2.31, *p* =.022, d = 0.17). We also observed a trend for a decrease of eccentricity from inferior-to-superior positions, which however did not reach significance (*M* = −0.03, *SD* = 0.21, *t*(180) = −1.87, *p* =.063, d = −0.14). We did not find a systematic eccentricity change along the left-right dimension in either the left (*M* = −0.10, *SD* = 1.33, *t*(167) = −0.96, *p* =.337, d = −0.07) or the right visual claustrum (*M* = 0.002, *SD* = 0.85, *t*(179) = 0.03, *p* =.973, d = 0.003).

**Figure 6.**
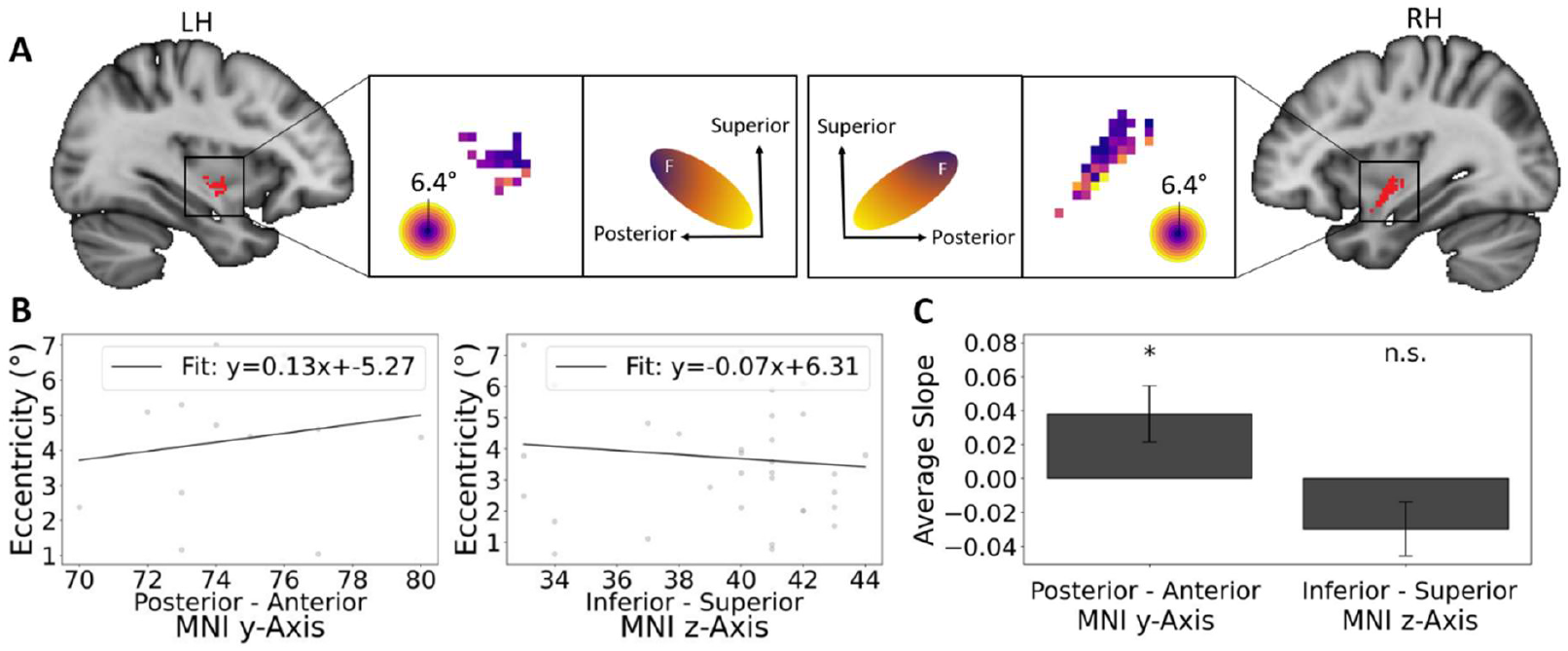
Relationship between the eccentricity representation and the brain axes. A) Eccentricity maps based on the averaged time courses across subjects (Sub-ID: 999999) shown on a sagittal cross-section of the left and right hemisphere. The middle parts show schematic illustrations of the inferred gradient of eccentricity representations for each hemisphere, extending from posterior-superior to anterior-inferior. B) Relationship between the eccentricity values and the posterior-anterior dimension (Sub-ID: 330324) and the inferior-superior dimension (Sub-ID: 200614) in example participants. C) Regression line slope values for the whole participant sample, demonstrating the significant nonzero increase along the posterior-anterior brain axis, and the non-significant trend for a decrease along the inferior-superior axis.

We did not detect any systematic change in polar angle values along the posterior-anterior dimension (*M* = 0.50, *SD* = 5.09, *t*(180) = 1.32, *p* =.190, d = 0.10), the inferior-superior dimension (*M* = −0.57, *SD* = 4.87, *t*(180) = −1.58, *p* =.117, d = −0.12), or the left-right axis for either the left (*M* = 2.47, *SD* = 30.93, *t*(167) = 1.04, *p* =.302, d = 0.08) or the right visual claustrum (*M* = −1.62, *SD* = 18.71, *t*(179) = −1.16, *p* =.246, d = −0.09).

## Discussion

The pRF mapping approach for functional Magnetic Resonance Imaging employed in this study allowed us to characterize visual field representation and pRF properties of the human visual claustrum zone. The claustrum exhibited properties typical for a visually responsive area such as the contralateral visual field bias and an increase in pRF size with increasing eccentricity. Compared to other low- and higher-level visual areas, the visual claustrum zone exhibited weaker lateralization of responses towards the contralateral hemifield and a weaker increase of receptive field sizes with increasing eccentricity. In contrast to other visual areas examined, it showed no horizontal meridian bias and a larger representation of the visual periphery. These results demonstrate the similarities of the claustrum to other low- and high-level visual areas but also reveal several properties that distinguish the claustrum. In the following, we discuss these aspects in the context of characteristics found in other species, speculating on the claustrum’s position in the visual processing hierarchy and its potential contribution to visual cognition.

The functional organization of the visual claustrum zone has so far been best characterized in felines and to a lesser extent in macaques. In felines, there is mixed evidence with regard to the topography of the visual zone, with some studies reporting no topographic organization (e.g., Jayaraman & Updyke, 1979), while others reported one retinotopically organized visual field map (Olson & Graybiel, 1980; LeVay & Sherk, 1981a,b). In the macaque (Remedios et al., 2010; Gattass et al., 2014), whose visual zone location is more similar to what has been reported in humans (Coates et al., 2024), a coarse retinotopic organization has been described (Gattass et al., 2014). Our results are broadly consistent with the macaque findings on the existence of a coarse topography. First, the human visual claustrum zone showed a less pronounced but significant bias towards the contralateral visual field. Second, a weak but significant progression in eccentricity values along the posterior-anterior and a non-significant trend along the inferior-superior dimension speak for the existence of at least a coarse retinotopic ordering. Altogether, these results confirm the existence of a coarse topographic representation of visual space in the human claustrum.

In this study we demonstrated an important property of the human visual claustrum zone that distinguishes it from most visually responsive regions. The visual claustrum contains a larger representation of more peripheral visual field regions compared to other early and late visual processing areas we examined. Previously, this topographic attribute of claustrum organization was only shown in felines (LeVay & Sherk, 1981b). The fact that this feature remained preserved in evolution up to the human species implies its ecological importance. In the visual cortex, such homogeneous representation of eccentricities is a key characteristic of motion area V6 (Galletti et al., 1999). A more homogeneous representation of all eccentricities, without emphasizing the fovea could be important for multiple functions such as optic flow processing, or reorienting attention to salient peripheral stimuli. The latter function is consistent with the claustrum’s proposed role in salience processing and distractor suppression based on macaque and rodent studies (Remedios et al., 2014; Atlan et al., 2018; Smith et al., 2019). A specific role of the human visual claustrum in reorienting attention to peripheral distractors or suppressing such a reorientation response should be tested in future studies.

Another interesting property that distinguishes the claustrum visual field representation is the apparent absence of the horizontal meridian bias. While the other examined regions exhibited a relative overrepresentation of the horizontal meridian, the visual claustrum zone exhibited a homogenous distribution of polar angles in all directions. The overrepresentation of the horizontal meridian is thought to stem from the distribution of photoreceptors in the retina, with heightened density of photoreceptors along the visual streak (Curcio et al, 1990). The fact that this bias is present in all examined cortical areas but not in the visual claustrum zone emphasizes the visual claustrum’s special position in the visual processing stream. Both feline and macaque studies demonstrated that the visual claustrum zone receives input from multiple cortical areas (Jayaraman & Updyke, 1979; Olson & Graybiel, 1980; LeVay & Sherk, 1981a; Gattass et al., 2014), but comparably less from subcortical structures (LeVay & Sherk, 1981a). Crucially, neither of the subcortical sensory nuclei has claustral projections (LeVay & Sherk, 1981a). Such a projection architecture together with the absence of a meridian bias defines the visual claustrum zone as a non-cortical higher-level stage of information processing, which is more abstracted from the sensory input-related biases present in the visual cortical areas.

Although the demonstrated presence of an overall coarsely ordered retinotopy is consistent with animal work, the specific layout of the topographic map inferred in our study diverges from a single previous report of the visual claustrum organization in macaques. Gattass and colleagues (2014) used tracer injections at different locations within visual area V4 to map the topographic organization of the claustrum, finding two distinct, coarse topographic maps, a smaller and less consistently found dorsal map and a larger ventral map. Within the larger ventral map, which is likely to dominate our fMRI results, the eccentricity values increase from inferior to superior positions, while in our data an opposite trend is observed. One potential reason for this discrepancy is that Gattass et al. (2014) focused on the topography of V4 projections only. It has been previously reported for the feline claustrum that projections from different visual and visuomotor cortical areas overlap within the visual claustrum zone (LeVay & Sherk, 1981b; Gattass et al., 2014). Although it is generally assumed that the topography of visual projections is preserved in the visual claustrum across different source regions in animals (LeVay & Sherk, 1981a,b; Gattass et al., 2014), nothing is known about the topography of the visual-to-claustrum projection system in humans. It is not clear whether different retinotopic areas consistently project to the same parts of the visual claustrum zone and which of the visual projections is the primary drive of the visual responses within the claustrum. Hence, the topography of projections from V4 may be inconsistent with our findings because projections from other areas follow a different organization principle. In fact, the significant gradient of eccentricities from posterior to anterior, which we observed in the claustrum, is consistent with the eccentricity gradient in the early visual cortex V1-V3, where values also increase from the occipital pole towards the anterior positions (Dumoulin & Wandell, 2008). This could mean that the projections of the early visual areas to the visual claustrum preserve the eccentricity ordering. Further invasive work in animals is needed to determine the topography of inputs to the visual claustrum from visual areas other than V4.

Despite its advantages, the pRF method has one limitation which hindered us in directly comparing the pRF size estimates of the claustrum with other examined areas. The estimated pRF size of a voxel is influenced by the true receptive field sizes of neurons in that voxel, the extent of visual space represented by those neurons (range of RF centers) and the amount of jitter of receptive field positions (Smith et al., 2001; Dumoulin & Wandell, 2008). Comparably small topographic maps like the visual claustrum zone will be inevitably biased towards larger pRF estimates compared to e.g. V1, just because a larger extend of visual space is represented by the same voxel volume in the former. Computational methods can - to some extent - overcome these confounds, but they are based on assumptions derived from the visual cortical processing and require special stimulus design (Keliris et al., 2019). Judging from the distribution of the pRF size estimates of the visual claustrum in comparison with LGN, which has a similar volume, it appears that visual claustrum neurons could have larger central receptive fields. However, due to the above-mentioned methodological problems, we emphasize that this must be confirmed by true neural receptive field size measurements in the primate visual claustrum zone. Knowledge of the receptive field properties of the claustrum neurons obtained using invasive electrophysiological recording methods is therefore critical for verifying pRF estimates of the visual claustrum zone in humans. As of today, such knowledge for the primate claustrum is missing.

### Conclusion

Leveraging the high sensitivity provided by 7 Tesla MRI and the high statistical power of the HCP dataset, we characterized the visual topographic organization of the human visual claustrum zone. Our findings demonstrate both characteristics common to typical visual cortical and subcortical areas, and features that appear to be unique to the claustrum as a higher-level visual processing area. This study is the first to characterize the topographic organization of the visual claustrum zone in humans, yielding important information for further investigation of the claustrum’s role in visual processing.

## Acknowledgments

This research was funded in whole or in part by the Austrian Science Fund (FWF) [Grant-DOI 1: 10.55776/PAT8722623; Grant-DOI 2: 10.55776/P35583]. For open access purposes, the author has applied a CC BY public copyright license to any author accepted manuscript version arising from this submission.

## Author contributions

C.P. and N.Z. designed research; C.P. and D.L. analyzed data; A.C. provided the visual claustrum mask; C.P. and N.Z. wrote the first paper draft and the paper; C.P., N.Z., D.L., A.C., D.Z. and C.W. edited the paper.

